# Mucociliary Clearance is Impaired in Small Airways of Cystic Fibrosis Pigs

**DOI:** 10.1101/2024.05.22.595427

**Authors:** Carley G. Stewart, Brieanna M. Hilkin, Nicholas D. Gansemer, David W. Dick, John J. Sunderland, David A. Stoltz, Mahmoud H. Abou Alaiwa, Joseph Zabner

## Abstract

**Rationale:** Cystic fibrosis is a genetic disorder characterized by recurrent airway infections, inflammation, and progressive decline in lung function. Autopsy and spirometry data suggest that cystic fibrosis may start in the small airways which, due to the fractal nature of the airways, account for most of the airway tree surface area. However, they are not easily accessible for testing.

**Objectives:** Here, we tested the hypothesis that mucociliary clearance is abnormal in the small airways of newborn cystic fibrosis pigs.

**Methods:** Current mucociliary clearance assays are limited therefore we developed a dynamic positron emission tomography scan assay with high spatial and temporal resolution. Each study was accompanied by a high-resolution computed tomography scan that helped identify the thin outer region of the lung that contained small airways.

**Measurements and Main Results:** Clearance of aerosolized [^68^Ga]macro aggregated albumin from distal airways occurred within minutes after delivery and followed a two-phase process. In cystic fibrosis pigs, both early and late clearance rates were slower. Stimulation of the cystic fibrosis airways with the purinergic agonist UTP further impaired late clearance. Only 1 cystic fibrosis pig treated with UTP out of 6 cleared more than 20% of the delivered dose.

**Conclusions:** These data indicate that mucociliary transport in the small airways is fast and can easily be missed if the acquisition is not fast enough. The data also indicate that mucociliary transport is impaired in small airways of cystic fibrosis pigs. This defect is exacerbated by stimulation of mucus secretions with purinergic agonists.

## INTRODUCTION

Cystic Fibrosis (CF) is a genetic disease caused by mutations in the *cystic fibrosis transmembrane conductance regulator* (*CFTR*) gene with high morbidity and mortality arising from airway involvement (1-3). Many observations suggest that the small airways are perhaps the site where the airway disease originates (4, 5). Old autopsy studies from young people who succumbed to CF and histopathology of lung explants from individuals with advanced CF airway disease getting lung transplant showed increased diameter of small airways, evidence of bronchiolitis obliterans, destructive emphysema, increased wall thickness, and decreased density of small airways (6). In children with CF, multiple breath washing (MBW) studies showed abnormalities in lung clearance index (LCI) (7). In adults with early CF airway disease, FEF_75_ and FEF_25-75_ are reduced and progress before any FEV_1_ or FVC defects are detected (8). Air trapping and ventilation defects are detected on computed tomography (CT) imaging and hyperpolarized ^3^He MRI scans very early in the disease course (9). The lack of a large animal model that recapitulates human CF airway disease and difficulties in accessing the small airways are reasons the hypothesis that CF originates in the small airways remained untested (2).

We developed a porcine animal model of CF with targeted disruption of the *CFTR* gene (10). Older CF pigs have the hallmarks of human CF airway disease including spontaneous bacterial infection, neutrophilic inflammation, and mucus plugging (10). Newborn CF pigs show no signs of infection or inflammation; yet their large airways are unable to kill bacteria or transport large particles (11). Using tantalum microdisks and dynamic CT scan imaging, we found that mucus strands secreted by submucosal glands are responsible for the CF MCT defect (12). In CF, the forward transport of microdisks is sometimes interrupted by mucus strands that fail to detach from submucosal gland ducts (12). In other situations, the forward transport is repeatedly reversed with abrupt backward recoil due to abnormally elastic mucus strands (13). Disrupting the mucus strand structure, with reducing agents, that break the disulfide bridges that hold the mucus together, reverse these defects (13). Since small airways lack submucosal glands, it is possible that MCT in the small airways is unaffected in CF.

The human lungs evolved with a unique fractal structure to achieve efficient gas exchange (14). The conducting airways divide sequentially to generate ∼2 daughter airways in a dichotomous branching pattern for ∼22 generations (15). After generation 15, the conducting airways transition into respiratory bronchioles and alveolar ducts with specialized epithelia for gas exchange (14). As the proximal airways divide, the total cross-sectional area of the airway tree increases exponentially (14). The small airways are the airways that contribute the least to total airway resistance (16). These airways have a distinct cuboidal appearance, lack submucosal glands, lack complete cartilage rings, and their secretory cells express surfactant protein D (SPD) (17). SPD has a role in reducing ASL surface tension and helps prevent airway closure on expiration (18). In young piglets, the airways with a diameter less the 200 *μ*m have these characteristics (19).

With each generation of branching, a parent airway splits into ∼2 smaller and shorter daughter airways (20, 21). With shorter distances to travel with each generation, mucus will be transported out faster in more distal airways. To capture and study this early clearance, we need an MCT assay with high temporal resolution and with instantaneous acquisition that can be started immediately after delivery of the particles. Because current MCC assays rely on pooling data for several minutes before clearance is calculated, the fast clearance may be missed. In addition, the planar nature of these assays allows overlap between the central and peripheral regions of interest. To circumvent these limitations, we developed a novel dynamic positron emission tomography (PET) technique to measure mucus transport with high spatial and temporal resolution in the small airways (22).

The tantalum microdisks that we used to study MCT in the large airways of pigs were too large to reach the small airways and much larger in comparison to physiologically relevant inhaled pathogens or particulates (12, 13, 23). Here, we used macro aggregated albumin (MAA) labelled with a positron emitting isotope and monitored the movement with dynamic PET scan imaging. Inspired by the use of Tc-99m macroaggregated albumin (MAA) to measure tracheal mucus velocity (24), we radiolabeled MAA with gallium-68. Gallium-68 was used because it is readily available, has a convenient short half-life of 68 min, decays into positrons of sufficient energy to create high quality images, easy to label with MAA (25), and facilitates the use of high temporal and spatial resolution, quantitative PET/CT rather than single photon imaging. Gallium-68 was attached to macroaggregated albumin with 15-30 *μ*m predicted particle size (26). The radiotracer was delivered via a catheter placed through the trachea and introduced into the distal airways and the data were acquired in list mode where the spatial location of each positron annihilation event was recorded, and time stamped. By acquiring the data in list mode, it can be reconstructed with different time scales ranging from a tenth of a second up to several minutes. On high resolution CT scans, we identified a thin outer shell of the lung that contained small airways, accumulated the activity in this region, and quantified clearance from the time-activity curve. Because small airways lack SMGs, we hypothesized that MCT in the small airways of newborn CF pigs would be intact.

## RESULTS

### Mucus clearance is impaired in the small airways of CF pigs

In prior reports, using scintigraphy techniques, clearance from distal lung regions was reported to occur in a time scale of hours rather than minutes (27). This reported slow clearance rate was likely due to the low temporal resolution of these imaging assays (28). To measure clearance from the small airways we first highlighted airways in a micro-CT reconstruction of the porpoise lobe (29). A shell with a depth of 1.5 mm enclosed most of the airways with a diameter less than 200 *μ*m (Fig. 1A). We then inscribed a 3-dimensional shallow shell within the outer lung and measured MCC (Fig. 1B). The radiotracer cleared out of this shell with time. Clearance started immediately within a minute after delivery (Fig. 1C). Early clearance rates were fast but slowed down after 5 min (Fig. 1C). In CF pigs, both early (non-CF 7.77±1.76 % cleared/min vs. CF 6.99±2.34%) and late (non-CF 0.64± 0.13% cleared/min vs. CF 0.36±0.08% cleared/min) clearance rates were slower. Purinergic stimulation results in mucus secretion, chloride secretion (TMEM), and increased ciliary beat frequency (30-32). However, stimulation of the CF airways with UTP further impaired late clearance (early 7.63±3.77% cleared/min, late 0.26%±0.16). Our results showed most non-CF pigs (57%, N=7) cleared more than 20% of the delivered dose within 15 minutes. Only 1 CF pig (8%, N=12) cleared more than 20% of the delivered dose during that time (Fig. 1D).

**Figure 1.**
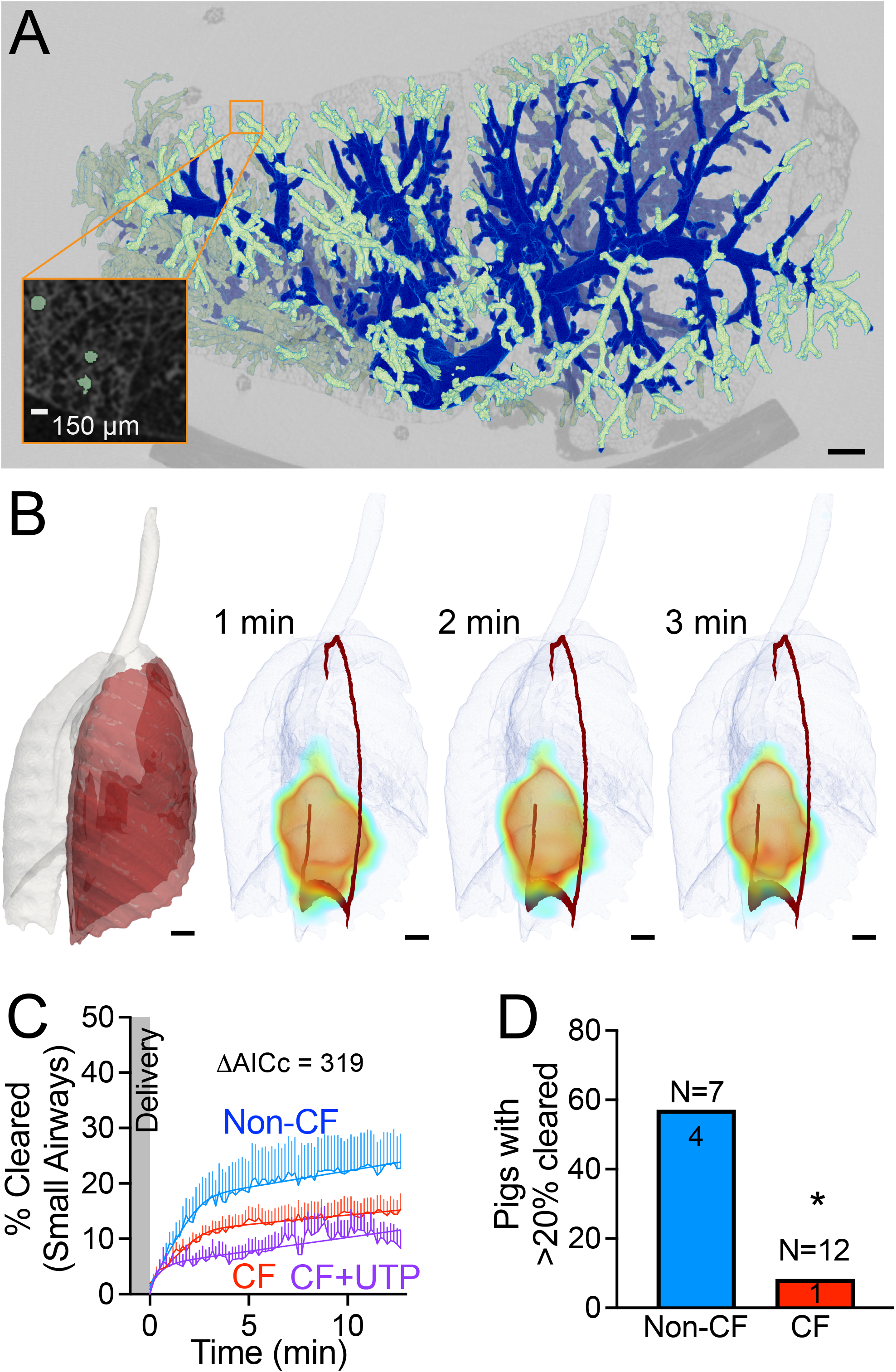
Mucociliary clearance is impaired in the small airways of CF pigs. **A**. Micro-CT of the porpoise lobe of newborn pig. Small airways (less than 200 *μ*m in diameter, light green) are present within a 1.5 mm shell inscribed in the outer lung (inset). Scale = 150 *μ*m. **B**. Images are composite PET signal superimposed on volumetric reconstruction of the lungs. Mucociliary is immediate and fast. Red represents cross section of a 1.5 mm shell. Scale = 10 mm. **C**. Percent change in the activity of delivered dose (%Cleared) from small airways as a function of time. Lines represent mean and standard error for non-CF (blue), CF (red) or CF with UTP (purple). Probability < 0.01% by AICc. **D**. Number of pigs with clearance greater than 20% at the end of the study. * P < 0.01 by Fisher’s exact test. Non-CF in blue and CF in red.

### Mucus transport is impaired in the distal airways of CF pigs

A concentric shell around the outer lung may include, in addition to the small airways, non-conducting airways such as respiratory bronchioles and alveolar spaces. Because clearance in non-conducting airways may be independent of mucus secretion or ciliary function (e.g. alveolar macrophage phagocytosis), we followed up with an alternate analysis approach. The high-resolution CT scan in our study of newborn pigs had limited resolving power to 2 mm in diameter airways. Therefore, we extended the centerline of the main lobar airway by extrapolation and sampled the radiotracer signal from a 2 mm diameter cylinder that extends to the visceral pleura (Fig. 2A). The radiotracer cleared from the distal airways quickly, within minutes of deposition (Fig. 2B). Similar to our findings in the previous technique, clearance from small airways followed a two-phase process, with an immediate brisk phase of clearance within the first 5 minutes after delivery followed by slower clearance (Fig. 2C). In CF pigs, both early (non-CF 12.88±8.04% cleared/min vs. CF 5.10±1.55% cleared/min) and late (non-CF 2.35±0.78% cleared/min vs. CF 0.33±0.1% cleared/min) clearance rates were slower. Stimulation of the CF airways with UTP further impaired late clearance (early 14.43 % cleared/min ± 7.9, late 0.18 % cleared/min ± 0.18) (Fig. 2C). As a result, several CF pigs (58.33 %, N=12) cleared more than 20% of the delivered dose. However, only 1 CF pig treated with UTP (16.67 %, N=6), cleared more than 20% of the delivered dose (Fig. 2D).

**Figure 2.**
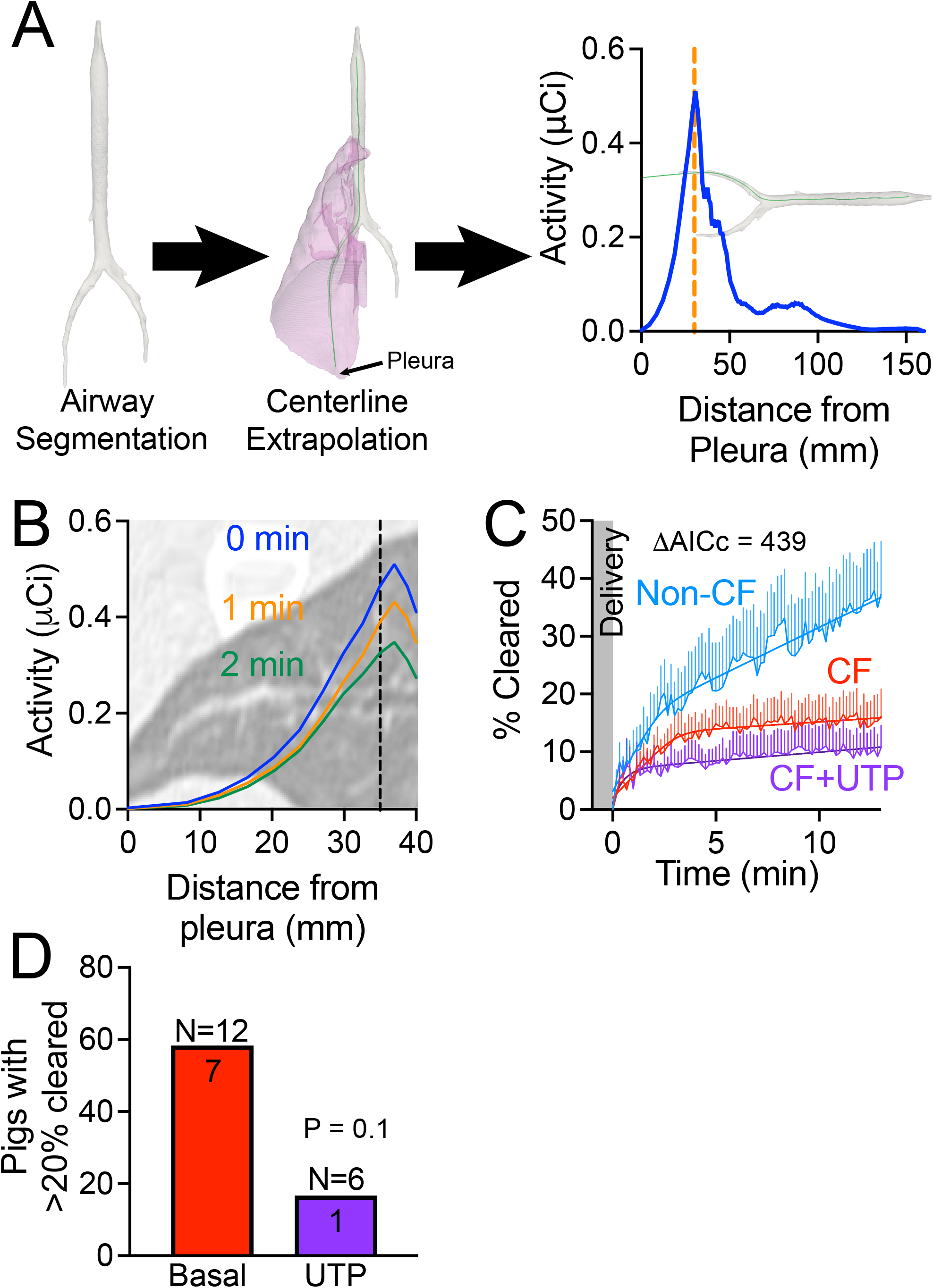
Mucociliary clearance is impaired in the distal airways of CF pigs. **A**. PET signal (activity in *μ*Ci) as we move away from the pleura. **B**. Clearance from distal airways is fast (in minutes). **C**. Percent change in the activity of delivered dose (%Cleared) from distal airways as a function of time. Lines represent mean and standard error for non-CF (blue), CF (red) or CF with UTP (purple). Probability < 0.01% by AICc. **D**. Number of CF pigs with clearance greater than 20% at the end of the study. CF in red and CF with UTP in purple.

## DISCUSSION

The submucosal gland duct opening is the site where mucus strands with abnormal biophysical properties get stuck and impede MCT in CF large airways (12, 13). The small airways lack submucosal glands (19) and thus MCT in CF small airways could have been normal. Our data indicate that MCT is also impaired in the small airways of newborn CF pigs. This defect is accentuated by stimulation of mucus secretion with purinergic agonists.

Mucus plugging or mucoviscidosis in small airways seen in autopsy studies is a common pathologic feature of CF (33). Inability to clear mucus that is secreted in the small airways may cause mucus accumulation and airway plugging (4, 34). Alternatively, some have suggested that the source for mucus plugs in the small airways is from mucus secreted in the large airways, traveling backwards (35). We described abrupt backward movement of abnormally elastic mucus strands secreted by submucosal glands (13). Perhaps, mucus strands snap, ricochet, and lodge in the small airways.

Our data support the former and provide an explanation for the origin of mucus plugs in the small airways. Inability to clear mucus in the small airways may cause mucus accumulation and airways plugging.

CF, like other pulmonary chronic airway inflammatory diseases, is characterized by goblet cell metaplasia and increased mucus secretion (36). In healthy small airways, the mucus gel is produced by columnar secretory cells that do not stain for intracellular mucins. The mucins are constitutively expressed in low amounts and are immediately secreted (37). Purinergic stimulation results in a marked increase in mucus secretion in goblet cells (>100 fold) (38) and also activates non-CFTR chloride channels, and increases ciliary beat frequency (32). In a mouse model of allergic goblet cell metaplasia induced by ovalbumin, ATP stimulation resulted in mucin airway occlusion (39).

In human CF small airways, most of the mucus plugs stain positive with Ab that recognizes MUC5B, even though MUC5AC is also seen (35). Secretory cells in small airways express mostly MUC5B, and goblet cells mostly MUC5AC (37). Because SMGs also secrete mostly MUC5B it is still possible that some of the mucus in small airways originates in SMGs (37, 40).

We previously reported that a combination of acidic ASL pH and decreased liquid secretion in the lumen of submucosal glands contributes to the abnormal viscoelastic properties of mucus strands (12, 13, 40). Small airway epithelial cultures obtained from pig CF airways have decreased Cl secretion, acidic ASL pH, and increased ASL viscosity (19). *In vivo* measurements are not available due to difficulties accessing the small airways. However, in flattened excised small airways, we found that CF airways cleared fluorescent microspheres slower than non-CF. Moreover, after purinergic stimulation most of the microspheres in CF pigs got stuck (41).

Inhalation deposition patterns of particles in the airways depends, in addition to physical and chemical characteristics of the particles, on the anatomy and physiology of the respiratory tract (42). The small airways of large mammals, including humans and pigs, are similar in size to the large airways of small animals such as mice. This distinction raises the question of whether, in small animals such as mice, the deposition of small particles is like that in large mammals such as pigs and humans and whether they are equipped with the same mucus clearance mechanisms. Interestingly, mice airways lack submucosal glands (43). In the large airways of humans and pigs with CF, CFTR-mediated HCO_3_^-^ secretion is lacking (44). As a result, H^+^ secretion by ATP12A is unopposed and the ASL is abnormally acidic (11). In mice with CF, the absence of CFTR has little effect on ASL pH because ATP12A is not expressed in mouse airways (45). This may be a reason why CF mice do not get CF airway disease. The small airways of humans and pigs do not express ATP12A either (45). They are more like mouse airways. Thus, the lack of CFTR-mediated HCO_3_^-^ secretion may have little effect on ASL pH and MCC. However, we found that pig small airways express a V-type proton pump ATPase (ATP6V0D) that is also expressed in the clear cells of the vas deferens (41). The localization of this pump on the cell luminal surface is pH-dependent (41). Under alkaline pH, it is translocated to the apical membrane where it results in net H^+^ secretion (41). The ASL pH in cultures of small airways from CF and non-CF pigs is alkaline enough (>7) to allow translocation of the V-ATPase to the apical membrane and, as a result, the CF small airway epithelial cultures were more acidic than their non-CF counterpart (11). Our data are consistent with an abnormal small airway MCT that may be related to abnormally acidic small airway ASL pH (41). However, these findings need to be confirmed with direct measurements *in vivo* and with interventions predicted to alkalinize the ASL.

Purinergic stimulation is predicted to induce Cl^-^ secretion from non-CFTR channels such as TMEM16 (46). Because of this, a P2Y receptor agonist was tested in people with CF. A phase III trial on 466 people with CF showed no clinical benefit of denufosol compared to placebo (47). Even though we have not measured ASL height in CF small airways, our data suggest that increased mucus secretion and the abnormal viscoelastic property of CF mucus outweigh the effects of increased ASL height on small airway MCT (48).

This study has many advantages. First, the PET/CT imaging system measures MCT *in vivo* with high temporal and spatial resolution (49, 50). Previously, the most common MCT assay captures clearance in two-dimensions with gamma-scintigraphy, while our setup, with high resolution CT (HRCT), allows for three-dimensional analysis (51). Gamma-scintigraphy methods typically measure retention of a radiotracer at 1–24-hour timepoints in concentric two-dimensional shells (52). Two-dimensional ROI’s make accurately assessing small and large airway contributions difficult, and hour-plus timepoints, although capturing non-ciliated clearance, miss the faster small airway clearance. In PET-CT, utilizing HRCT images, we can analyze anatomically positioned ROIs in three-dimensions. A ring of PET detectors and positron emitting isotopes, in this case gallium-68, allow for instantaneous data acquisition. Each event is timestamped, recorded, and can be reconstructed into images with time frames on the order of seconds capturing dynamic fast clearance (49). Second, we studied a newborn spontaneously breathing porcine model. Pigs have gland-containing proximal airways, much like humans, and the CF porcine model reproduces a similar phenotype to that seen in people with CF (2). Additionally, newborn CF pigs lack infection, inflammation, and airway remodeling allowing us solely to study the host defense defects at an early timepoint (11, 53). Third, we delivered radiotracer particles of relevant size directly into the distal airways. Typical assays deposit radioaerosol through specific inhalation techniques, which results in some tracer depositing centrally even when targeting peripheral airways. A distal delivery ensures deposition in the most distal airways.

This work also has limitations. First, HRCT imaging could not resolve small airways defined in newborn pigs as <200 μm (19). To circumvent this limitation, we analyzed two small airway regions. A traditional concentric shell with a depth found to encompass small airways and a region extending from the main airway to the pleura. Second, these regions are limited. The small airway shell will contain alveoli, which will retain deposited radiotracer until removal by macrophages. This most likely causes an underestimation of clearance due to background spillover of the alveoli deposited radiotracer. The extended region to the pleura starts from an airway with a diameter of 2 mm. This is likely to encompass some airways with submucosal glands. Third, lung disease is known to be heterogenous, and here only a subset of small airways is analyzed.

This work, along with previous studies, further characterizes mucus transport in small and large airways in healthy and CF pigs (12, 13, 23, 40, 54). The fast temporal aspect of PET allowed us to assess MCT in small airways at relevant time points. Therapeutic interventions aimed at treating small airway disease in CF will benefit from this technique. Moreover, since small airways have been implicated in the pathogenesis of diseases that lack large animal models like asthma, COPD, and MUC5B promoter variant rs35705950 related IPF, it would be important to translate this methodology to study humans with these diseases (55, 56).

## METHODS

### Animals

We previously reported production of CFTR^-/-^ pigs (53). Newborn pigs were obtained from Exemplar Genetics. Animals were studied 8-15 hours post-birth. Sedation was with ketamine (20 mg/kg, intramuscular [IM]; Phoenix Pharmaceutical, Inc.) and acepromazine (2mg/kg, IM; Phoenix Pharmaceutical, Inc.) and anesthesia was maintained with IV dexmedetomidine (10 mg/kg/h; Accord Healthcare, Inc.). Euthanasia was with IV Euthasol (Virbac). The University of Iowa Animal Care and Use Committee approved all animal studies.

### Micro-CT

The tracheal lobe was scanned at 20 cm H_2_O with a Siemens micro-CAT II scanner (Siemens, Pre-Clinical Solutions; Knoxville, TN) and reconstructed with an isotropic voxel spacing of 0.028 mm.

### *In vivo* MCT assay

To measure MCT in small and large airways, we used a previously described PET based MCT assay (22). Gallium-68 MAA was obtained by eluting a germanium-68/gallium-68 generator and applying a commercial MAA labeling kit (Pulmotech MAA; Curium US, Maryland Heights, MO). [Ga-68] MAA particles (∼15-30 *μ*m, 15-100 mCi) were delivered using 1.5 mm polyethylene tubing (Intramedic; Parsippany, NJ) introduced into the distal airways of sedated newborn pigs. We acquired PET data in a continuous list mode for 15 min. We also acquired high resolution CT scans for anatomical references. We binned the data into 10 second timeframes to reconstruct the PET images.

### Analysis

We quantified the PET signal in the distal airway region by either extending the airway volume to the pleura or constructing a region of interest in a uniform thin shell.

### Statistical analysis

Differences were considered statistically significant at P < 0.01. All analyses completed in GraphPad Prism v9.5.1 (GraphPad Software, La Jolla, CA). Data are presented as mean ± SEM are indicated by bars. For %Cleared in Fig. 1C and Fig. 2C, we modeled slow and fast clearance phases with a 4-parameters continuous hinge function (segmental linear regression by least square method). We used Akaike’s Information Criterion corrected for small samples (AICc) to reject the simpler model that all 4-parameters are the same for all data. The alternative model states that all 4 parameters are different for at least one data set. We calculated the probability that a model is correct using the change (ΔAICc) in Akaike’s Information Criterion corrected for small samples with the following formula: 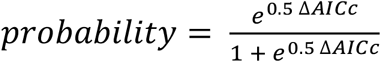 . We used Fisher’s exact test to calculate p-value for contingency data in Fig. 1D and Fig. 2D.

### Study approval

The present studies in animals were reviewed and approved by the University of Iowa Animal Care and Use Committee.

## Data Availability

All study data are included in the article and/or supporting information. Code for PET analysis is available from the corresponding author upon request.

## ACKNOWLEDGEMENTS

We thank Jess Jensen for help with the manuscript preparation. We thank the Small Animal Imaging Core Facility and the PET Center at the Department of Radiology at the University of Iowa.

## REFERENCES

1. Ratjen F, Bell SC, Rowe SM, Goss CH, Quittner AL, Bush A. Cystic fibrosis. Nature Reviews Disease Primers 2015; 1: 15010.

2. Stoltz DA, Meyerholz DK, Welsh MJ. Origins of cystic fibrosis lung disease. The New England journal of medicine 2015; 372: 1574–1575.

3. Cutting GR. Cystic fibrosis genetics: from molecular understanding to clinical application. Nat Rev Genet 2015; 16: 45–56.

4. Ratjen F. Cystic fibrosis: the role of the small airways. Journal of aerosol medicine and pulmonary drug delivery 2012; 25: 261–264.

5. Tiddens HA, Donaldson SH, Rosenfeld M, Pare PD. Cystic fibrosis lung disease starts in the small airways: can we treat it more effectively? Pediatr Pulmonol 2010; 45: 107–117.

6. Ornoy A, Arnon J, Katznelson D, Granat M, Caspi B, Chemke J. Pathological confirmation of cystic fibrosis in the fetus following prenatal diagnosis. Am J Med Genet 1987; 28: 935–947.

7. Gustafsson PM, De Jong PA, Tiddens HA, Lindblad A. Multiple-breath inert gas washout and spirometry versus structural lung disease in cystic fibrosis. Thorax 2008; 63: 129–134.

8. Bakker EM, Borsboom GJ, van der Wiel-Kooij EC, Caudri D, Rosenfeld M, Tiddens HA. Small airway involvement in cystic fibrosis lung disease: routine spirometry as an early and sensitive marker. Pediatr Pulmonol 2013; 48: 1081–1088.

9. Robinson TE, Goris ML, Zhu HJ, Chen X, Bhise P, Sheikh F, Moss RB. Dornase alfa reduces air trapping in children with mild cystic fibrosis lung disease: a quantitative analysis. Chest 2005; 128: 2327–2335.

10. Stoltz DA, Meyerholz DK, Pezzulo AA, Ramachandran S, Rogan MP, Davis GJ, Hanfland RA, Wohlford-Lenane C, Dohrn CL, Bartlett JA, Nelson GAt, Chang EH, Taft PJ, Ludwig PS, Estin M, Hornick EE, Launspach JL, Samuel M, Rokhlina T, Karp PH, Ostedgaard LS, Uc A, Starner TD, Horswill AR, Brogden KA, Prather RS, Richter SS, Shilyansky J, McCray PB, Jr., Zabner J, Welsh MJ. Cystic fibrosis pigs develop lung disease and exhibit defective bacterial eradication at birth. Science translational medicine 2010; 2: 29ra31.

11. Pezzulo AA, Tang XX, Hoegger MJ, Alaiwa MH, Ramachandran S, Moninger TO, Karp PH, Wohlford-Lenane CL, Haagsman HP, van Eijk M, Banfi B, Horswill AR, Stoltz DA, McCray PB, Jr., Welsh MJ, Zabner J. Reduced airway surface pH impairs bacterial killing in the porcine cystic fibrosis lung. Nature 2012; 487: 109–113.

12. Hoegger MJ, Fischer AJ, McMenimen JD, Ostedgaard LS, Tucker AJ, Awadalla MA, Moninger TO, Michalski AS, Hoffman EA, Zabner J, Stoltz DA, Welsh MJ. Impaired mucus detachment disrupts mucociliary transport in a piglet model of cystic fibrosis. Science 2014; 345: 818–822.

13. Pino-Argumedo MI, Fischer AJ, Hilkin BM, Gansemer ND, Allen PD, Hoffman EA, Stoltz DA, Welsh MJ, Abou Alaiwa MH. Elastic mucus strands impair mucociliary clearance in cystic fibrosis pigs. Proceedings of the National Academy of Sciences of the United States of America 2022; 119: e2121731119.

14. Weibel ER. Lung morphometry: the link between structure and function. Cell Tissue Res 2017; 367: 413–426.

15. Horsfield K, Cumming G. Morphology of the bronchial tree in man. J Appl Physiol 1968; 24: 373–383.

16. Macklem PT. The physiology of small airways. American journal of respiratory and critical care medicine 1998; 157: S181–183.

17. Thurman AL, Li X, Villacreses R, Yu W, Gong H, Mather SE, Romano-Ibarra GS, Meyerholz DK, Stoltz DA, Welsh MJ, Thornell IM, Zabner J, Pezzulo AA. A Single-Cell Atlas of Large and Small Airways at Birth in a Porcine Model of Cystic Fibrosis. American journal of respiratory cell and molecular biology 2022; 66: 612–622.

18. Crouch E, Persson A, Chang D, Heuser J. Molecular structure of pulmonary surfactant protein D (SP-D). The Journal of biological chemistry 1994; 269: 17311–17319.

19. Li X, Tang XX, Vargas Buonfiglio LG, Comellas AP, Thornell IM, Ramachandran S, Karp PH, Taft PJ, Sheets K, Abou Alaiwa MH, Welsh MJ, Meyerholz DK, Stoltz DA, Zabner J. Electrolyte transport properties in distal small airways from cystic fibrosis pigs with implications for host defense. American journal of physiology Lung cellular and molecular physiology 2016; 310: L670–679.

20. Weibel ER. What makes a good lung? Swiss Med Wkly 2009; 139: 375–386.

21. Nelson TR, West BJ, Goldberger AL. The fractal lung: universal and species-related scaling patterns. Experientia 1990; 46: 251–254.

22. Stewart CG, Hilkin B, Gansemer ND, Walsh SA, Acevado MR, Akurathi V, Pandya DN, Comellas AF, Wadas TJ, Dick DW, Sunderland JJ, Stoltz DA, Welsh MJ, Abou Alaiwa MH. Measurement of Mucociliary Transport: Novel Application of Positron Emission Tomography. 2022 Ieee International Symposium on Biomedical Imaging (Ieee Isbi 2022) 2022.

23. Fischer AJ, Pino-Argumedo MI, Hilkin BM, Shanrock CR, Gansemer ND, Chaly AL, Zarei K, Allen PD, Ostedgaard LS, Hoffman EA, Stoltz DA, Welsh MJ, Alaiwa MHA. Mucus strands from submucosal glands initiate mucociliary transport of large particles. JCI Insight 2019; 4.

24. Morgan L, Pearson M, de Iongh R, Mackey D, van der Wall H, Peters M, Rutland J. Scintigraphic measurement of tracheal mucus velocity in vivo. Eur Respir J 2004; 23: 518–522.

25. Martiniova L, De Palatis L, Etchebehere E, Ravizzini G. Gallium-68 in Medical Imaging. Current Radiopharmaceuticals 2016; 9: 187–207.

26. Mathias CJ, Green MA. A convenient route to [68Ga]Ga-MAA for use as a particulate PET perfusion tracer. Appl Radiat Isot 2008; 66: 1910–1912.

27. Daviskas E, Anderson SD, Gonda I, Chan HK, Cook P, Fulton R. Changes in mucociliary clearance during and after isocapnic hyperventilation in asthmatic and healthy subjects. Eur Respir J 1995; 8: 742–751.

28. Agnew JE, Pavia D, Clarke SW. Airways penetration of inhaled radioaerosol: an index to small airways function? European journal of respiratory diseases 1981; 62: 239–255.

29. Bauer C, Adam R, Stoltz DA, Beichel RR. Computer-aided analysis of airway trees in micro-CT scans of ex vivo porcine lung tissue. Comput Med Imaging Graph 2012; 36: 601–609.

30. Lazarowski ER, Boucher RC. Purinergic receptors in airway epithelia. Curr Opin Pharmacol 2009; 9: 262–267.

31. Delmotte P, Sanderson MJ. Ciliary beat frequency is maintained at a maximal rate in the small airways of mouse lung slices. American journal of respiratory cell and molecular biology 2006; 35: 110–117.

32. Namkung W, Yao Z, Finkbeiner WE, Verkman AS. Small-molecule activators of TMEM16A, a calcium-activated chloride channel, stimulate epithelial chloride secretion and intestinal contraction. FASEB journal : official publication of the Federation of American Societies for Experimental Biology 2011; 25: 4048–4062.

33. Dickey LB. The Pulmonary Aspects of Cystic Fibrosis of the Pancreas. Cal West Med 1942; 57: 41–42.

34. Quinton PM. Both Ways at Once: Keeping Small Airways Clean. Physiology (Bethesda) 2017; 32: 380–390.

35. Burgel PR, Montani D, Danel C, Dusser DJ, Nadel JA. A morphometric study of mucins and small airway plugging in cystic fibrosis. Thorax 2007; 62: 153–161.

36. Ma J, Rubin BK, Voynow JA. Mucins, Mucus, and Goblet Cells. Chest 2018; 154: 169–176.

37. Okuda K, Chen G, Subramani DB, Wolf M, Gilmore RC, Kato T, Radicioni G, Kesimer M, Chua M, Dang H, Livraghi-Butrico A, Ehre C, Doerschuk CM, Randell SH, Matsui H, Nagase T, O’Neal WK, Boucher RC. Localization of Secretory Mucins MUC5AC and MUC5B in Normal/Healthy Human Airways. American journal of respiratory and critical care medicine 2019; 199: 715–727.

38. Jaramillo AM, Azzegagh Z, Tuvim MJ, Dickey BF. Airway Mucin Secretion. Ann Am Thorac Soc 2018; 15: S164–S170.

39. Morgan LE, Jaramillo AM, Shenoy SK, Raclawska D, Emezienna NA, Richardson VL, Hara N, Harder AQ, NeeDell JC, Hennessy CE, El-Batal HM, Magin CM, Grove Villalon DE, Duncan G, Hanes JS, Suk JS, Thornton DJ, Holguin F, Janssen WJ, Thelin WR, Evans CM. Disulfide disruption reverses mucus dysfunction in allergic airway disease. Nat Commun 2021; 12: 249.

40. Ostedgaard LS, Moninger TO, McMenimen JD, Sawin NM, Parker CP, Thornell IM, Powers LS, Gansemer ND, Bouzek DC, Cook DP, Meyerholz DK, Abou Alaiwa MH, Stoltz DA, Welsh MJ. Gel-forming mucins form distinct morphologic structures in airways. Proceedings of the National Academy of Sciences of the United States of America 2017; 114: 6842–6847.

41. Li X, Villacreses R, Thornell IM, Noriega J, Mather S, Brommel CM, Lu L, Zabner A, Ehler A, Meyerholz DK, Stoltz DA, Zabner J. V-Type ATPase Mediates Airway Surface Liquid Acidification in Pig Small Airway Epithelial Cells. American journal of respiratory cell and molecular biology 2021; 65: 146–156.

42. Schlesinger RB. Comparative deposition of inhaled aerosols in experimental animals and humans: a review. J Toxicol Environ Health 1985; 15: 197–214.

43. Wine JJ, Joo NS. Submucosal glands and airway defense. Proceedings of the American Thoracic Society 2004; 1: 47–53.

44. Smith JJ, Welsh MJ. cAMP stimulates bicarbonate secretion across normal, but not cystic fibrosis airway epithelia. The Journal of clinical investigation 1992; 89: 1148–1153.

45. Shah VS, Meyerholz DK, Tang XX, Reznikov L, Abou Alaiwa M, Ernst SE, Karp PH, Wohlford-Lenane CL, Heilmann KP, Leidinger MR, Allen PD, Zabner J, McCray PB, Jr., Ostedgaard LS, Stoltz DA, Randak CO, Welsh MJ. Airway acidification initiates host defense abnormalities in cystic fibrosis mice. Science 2016; 351: 503–507.

46. Knowles MR, Clarke LL, Boucher RC. Activation by extracellular nucleotides of chloride secretion in the airway epithelia of patients with cystic fibrosis. The New England journal of medicine 1991; 325: 533–538.

47. Ratjen F, Durham T, Navratil T, Schaberg A, Accurso FJ, Wainwright C, Barnes M, Moss RB, Group T-SI. Long term effects of denufosol tetrasodium in patients with cystic fibrosis. Journal of cystic fibrosis : official journal of the European Cystic Fibrosis Society 2012; 11: 539–549.

48. Kellerman D, Evans R, Mathews D, Shaffer C. Inhaled P2Y2 receptor agonists as a treatment for patients with Cystic Fibrosis lung disease. Adv Drug Deliv Rev 2002; 54: 1463–1474.

49. Snyder DL. Parameter-Estimation for Dynamic Studies in Emission-Tomography Systems Having List-Mode Data. Ieee Transactions on Nuclear Science 1984; 31: 925–931.

50. Guobao W. High Temporal-Resolution Dynamic PET Image Reconstruction Using a New Spatiotemporal Kernel Method. IEEE Trans Med Imaging 2019; 38: 664–674.

51. Olseni L, Wollmer P. Mucociliary clearance in healthy men at rest and during exercise. Clin Physiol 1990; 10: 381–387.

52. Langenback EG, Bergofsky EH, Halpern JG, Foster WM. Supramicron-sized particle clearance from alveoli: route and kinetics. Journal of applied physiology 1990; 69: 1302–1308.

53. Rogers CS, Stoltz DA, Meyerholz DK, Ostedgaard LS, Rokhlina T, Taft PJ, Rogan MP, Pezzulo AA, Karp PH, Itani OA, Kabel AC, Wohlford-Lenane CL, Davis GJ, Hanfland RA, Smith TL, Samuel M, Wax D, Murphy CN, Rieke A, Whitworth K, Uc A, Starner TD, Brogden KA, Shilyansky J, McCray PB, Jr., Zabner J, Prather RS, Welsh MJ. Disruption of the CFTR gene produces a model of cystic fibrosis in newborn pigs. Science 2008; 321: 1837–1841.

54. Ash JJ, Hilkin BM, Gansemer ND, Hoffman EA, Zabner J, Stoltz DA, Abou Alaiwa MH. Tromethamine improves mucociliary clearance in cystic fibrosis pigs. Physiol Rep 2022; 10: e15340.

55. Usmani OS, Han MK, Kaminsky DA, Hogg J, Hjoberg J, Patel N, Hardin M, Keen C, Rennard S, Ble FX, Brown MN. Seven Pillars of Small Airways Disease in Asthma and COPD: Supporting Opportunities for Novel Therapies. Chest 2021; 160: 114–134.

56. Seibold MA, Wise AL, Speer MC, Steele MP, Brown KK, Loyd JE, Fingerlin TE, Zhang W, Gudmundsson G, Groshong SD, Evans CM, Garantziotis S, Adler KB, Dickey BF, du Bois RM, Yang IV, Herron A, Kervitsky D, Talbert JL, Markin C, Park J, Crews AL, Slifer SH, Auerbach S, Roy MG, Lin J, Hennessy CE, Schwarz MI, Schwartz DA. A common MUC5B promoter polymorphism and pulmonary fibrosis. The New England journal of medicine 2011; 364: 1503–1512.

